# Role of *Dusp*5 KO on Vascular Properties of Middle Cerebral Artery in Rats

**DOI:** 10.1101/2023.12.04.569939

**Authors:** Chengyun Tang, Huawei Zhang, Jane J. Border, Yedan Liu, Xing Fang, Joshua R. Jefferson, Andrew Gregory, Claire Johnson, Tae Jin Lee, Shan Bai, Ashok Sharma, Seung Min Shin, Hongwei Yu, Richard J. Roman, Fan Fan

**Author notes:** Send Correspondence to: Fan Fan, M.D., M.S., FAHA Department of Physiology Medical College of Georgia Augusta University 1462 Laney Walker Blvd Augusta, GA 30912. These authors contributed equally to this work (Chengyun Tang, Huawei Zhang, Jane J. Border).

## Abstract

Vascular aging influences hemodynamics, elevating risks for vascular diseases and dementia. We recently demonstrated that knockout (KO) of *Dusp5* enhances cerebral and renal hemodynamics and cognitive function. This improvement correlates with elevated pPKC and pERK1/2 levels in the brain and kidneys. Additionally, we observed that *Dusp5* KO modulates the passive mechanical properties of cerebral and renal arterioles, associated with increased myogenic tone at low pressure, enhanced distensibility, greater compliance, and reduced stiffness. The present study evaluates the structural and mechanical properties of the middle cerebral artery (MCA) in *Dusp5* KO rats. We found that vascular smooth muscle cell layers and the collagen content in the MCA wall are comparable between *Dusp5* KO and control rats. The internal elastic lamina in the MCA of *Dusp5* KO rats exhibits increased thickness, higher autofluorescence intensity, smaller fenestrae areas, and fewer fenestrations. Despite an enhanced myogenic response and tone of the MCA in *Dusp5* KO rats, other passive mechanical properties, such as wall thickness, cross-sectional area, wall-to-lumen ratio, distensibility, incremental elasticity, circumferential wall stress, and elastic modulus, do not significantly differ between strains. These findings suggest that while *Dusp5* KO has a limited impact on altering the structural and mechanical properties of MCA, its primary role in ameliorating hemodynamics and cognitive functions is likely attributable to its enzymatic activity on cerebral arterioles. Further research is needed to elucidate the specific enzymatic mechanisms and explore potential clinical applications in the context of vascular aging.

## INTRODUCTION

Vascular aging denotes age-related structural and functional changes in blood vessels. These mechanical alterations in the vasculature influence hemodynamics, thereby heightening the risk of vascular diseases and dementia.^1^ Genetic factors, alongside various other mechanisms such as inflammation, mitochondrial dysfunction, oxidative stress, and telomere attrition, wield considerable influence over vascular aging by modulating vascular structure and hemodynamics. ^2–9^

Dual-specificity protein phosphatase 5 (DUSP5) is a nuclear enzyme that specifically dephosphorylates the tyrosine-threonine residues of extracellular signal-related kinase 1/2 (ERK 1/2) in the mitogen-activated protein kinase pathway, leading to the inactivation of ERK 1/2. ^10–13^ Elevations in phosphorylated protein kinase C (pPKC) and phosphorylated ERK1/2 (pERK1/2) promote calcium influx in vascular smooth muscle cells (VSMCs) and facilitate vasoconstriction. Knockout (KO) of *Dusp*5 results in ERK nuclear translocation, which promotes cell proliferation through heightened activities of cell cycle regulatory proteins and posttranslational modifications that enhance pro-survival gene function and reduce cell death.^11, 14^ DUSP5 also plays a direct role in promoting cell death in response to endoplasmic reticulum stress. ^15^ DUSP5 suppresses the actin cytoskeleton rearrangement by negatively modulating the ERK-MLCK-Myosin IIB signaling cascades for its potential antiviral effects. ^16^ The essential impacts of DUSP5 have been linked to ischemic stroke, hypertension, renal disease, pulmonary hypertension, cancers, osteoarthritis, immunological diseases, and neurodegenerative diseases. ^17–23^ In human studies, DUSP5 has been reported to regulate hippocampal dentate gyrus plasticity potentially and is associated with late-onset Alzheimer’s Disease (AD). ^24–26^

There is growing evidence suggesting that vascular factors can contribute to the development and progression of Alzheimer’s Disease and Alzheimer’s Disease-Related Dementias (AD/ADRD).^27, 28^ Brain hypoperfusion attributed to impaired cerebral blood flow (CBF) autoregulation, neurovascular uncoupling, blood-brain barrier leakage, and neurodegeneration are important potential causal factors in AD and aging-, hypertension-, and diabetes-related ADRD. ^4, 29–35^ We have recently documented that *Dusp5* KO amplifies myogenic reactivity and autoregulation of blood flow in both cerebral and renal circulations, concomitant with elevated levels of pERK1/2 and pPKC. ^10, 11, 22, 36, 37^ Inhibition of ERK and PKC notably boosted the contractile potential of VSMCs obtained from middle cerebral arteries (MCAs), associated with a dose-dependent dilation of the MCAs and penetrating arterioles (PAs), with a more pronounced impact observed in *Dusp5* KO rats. ^11, 36^ Additionally, there was a noteworthy enhancement in learning and memory observed in *Dusp5* KO rats, suggesting that targeting DUSP5 deletion could hold promise as a therapeutic approach for delaying vascular aging and the onset of AD/ADRD.^36^ Furthermore, we examined the impact of *Dusp5* KO on the mechanical characteristics of brain PAs and renal interlobular arterioles. We found that both PAs and renal IAs in *Dusp5* KO rats displayed features of eutrophic vascular hypotrophy, increased myogenic tone, improved distensibility, enhanced compliance, and reduced stiffness, while wall-to-lumen ratios remained unchanged. ^11^

The present study focused on how the KO of Dusp5 influenced the mechanical vascular characteristics of the MCA, the major blood vessel in the brain susceptible to various vascular disorders. ^38, 39^ Our study involved a thorough comparison of multiple factors in the MCA between *Dusp5* KO and wild-type (WT) rats. This comprehensive analysis encompassed the intrinsic structural characteristics, myogenic reactivity across a broad range of perfusion pressures (ranging from 40 to 180 mmHg), evaluation of myogenic tone, and assessment of passive mechanical properties.

## MATERIALS AND METHODS

### Animals

All experiments were conducted on 9- to 12-week-old male WT (FHH.1^BN-(*D1Rat*09*-*^ *^D1Rat^*^225)/Mcwi^) and *Dusp5* KO (FHH-Chr 1^BN^-*Dusp5*^em1Mcwi^) rats that we previously generated and robustly validated.^10, 11, 22, 36^ The rats were accommodated in the animal facility at the University of Mississippi Medical Center (UMMC), exposed to a 12-hour light-dark cycle with unrestricted access to a standard diet (Teklad rodent diet 8604, Envigo, Indianapolis, IN) and *ad libitum* water. All procedures conducted in this study were approved by the Institutional Animal Care and Use Committee at UMMC and were meticulously designed and executed in full compliance with the guidelines established by the American Association for the Accreditation of Laboratory Animal Care.

### Assessment of MCA Intrinsic Structural Features

#### Sample Preparation

Samples of the MCA were prepared in accordance with protocols that had been previously published. ^39^ Briefly, rats were perfused with 10% neutral-buffered formalin (Sigma-Aldrich, St. Louis, MO) intracardially at 100 mmHg after being anesthetized with 2% isoflurane. The brains were retrieved and immersed in 10% formalin for a duration of two days for fixation. An approximately 5 mm^3^ rectangular piece of brain tissue containing the M1 and M2 segments of the MCA was carefully dissected and embedded with paraffin. Serial cross-sections with a thickness of 3 μm were obtained, encompassing the initial 1 to 2 mm of the M2 segment of the MCA. This segment of the MCA exhibits a consistently cylindrical shape, characterized by vessels with a uniform cylindrical morphology. A systematic random sampling approach was utilized for stereological analysis. This involved the random selection of the first section from among the initial 10 sections, followed by the inclusion of every 10^th^ section thereafter. Four to six sections were evaluated in each animal in this study.

#### Assessment of VSMC Content in the MCA

Evaluation of VSMC content in the MCA wall was conducted through immunofluorescent staining with α-smooth muscle actin (α-SMA) antibody following our published protocol.^39^ The formalin-fixed and paraffin-embedded MCA sections underwent deparaffinization using xylene. Subsequently, xylene was removed, and the sections were rehydrated with 100%, 95%, and 70% ethanol, and tap water. After antigen retrieval with proteinase K (S3020, Agilent, Santa Clara, CA), the sections were blocked using Protein Block Serum-Free Blocking (X0909, Agilent, Santa Clara) and incubated with a primary antibody targeting mouse α-SMA (1:300, A5228, Sigma-Aldrich), followed by a secondary antibody, goat anti-mouse (1:1000, A21424, Thermo Fisher Scientific). An anti-fade mounting medium with DAPI (H-1200, Vector Laboratories, Burlingame, CA) was applied, followed by the placement of coverslips. Images were obtained with a Nikon C2 laser scanning confocal system on an Eclipse Ti2 inverted microscope (Nikon, Melville, NY) using a 60X oil immersion objective lens and applying a 3X digital zoom, resulting in a total magnification of 3,200X. With a z-step of 1.15 μm, a series of optical sections were captured, and a z-projection was created using three images. The evaluation of α-SMA expression involved comparing fluorescence average intensities per cross-sectional area (CSA) using NIS-Elements Imaging Software 4.6. Furthermore, VSMC counts were manually conducted by visually observing the distinct intercellular spaces between each cell relative to the CSA.

#### Assessment of Collagen Content in the MCA

Evaluation of collagen content within the MCA wall involved the use of Masson’s trichrome staining, and images were captured using a Nikon Eclipse 55i microscope and a DS-FiL 1 color camera. Collagen quantification within the media and adventitial layers of the MCA wall was achieved by comparing the average intensities of the blue staining with NIS-Elements Imaging Software 4.6.

#### Assessment of Elastin Content in the MCA

Evaluation of elastin content in the MCA wall relied on the observation that a linear relationship exists between autofluorescence intensity and elastin content, ^40, 41^ following a protocol we previously published. ^39^ Briefly, freshly isolated M2 segments of MCA were fixed with 4% paraformaldehyde at 100 mmHg at 37°C for 1 hour after being cannulated onto glass pipettes in a pressure myograph chamber (Living System Instrumentation, Burlington, VT) and equilibrated in calcium-free physiological salt solution (PSS_0Ca_). Non-elastin components were removed by incubating with 0.1 M sodium hydroxide at 75 °C for 1 hour. ^39, 40^ Following the removal from the pressure myograph, the vessels were placed onto slides containing a silicon spacer (500 μm depth, 13 mm diameter; Grace Bio Labs, Bend, Oregon) and immersed in VECTASHIELD anti-fade mounting medium (H-1000, Vector Laboratories). Elastin autofluorescence was compared based on the mean autofluorescence intensities per view, which were measured using NIS-Elements Imaging Software 4.6. These measurements were taken from images of multiple optical sections captured at wavelengths of excitation/emission 488/500–560 nm, utilizing a Nikon C2 laser scanning confocal system on an Eclipse Ti2 inverted microscope with a total magnification of 3,200X, as described above. Furthermore, comparisons were made for the thickness of the internal elastic lamina (IEL), the number of fenestrae, and the fenestrae area. ^40, 42, 43^

### Pressure Myography

#### Isolation and Preparation of MCA

The MCAs were isolated and prepared following the protocol we had previously documented. ^2, 6, 44, 45^ In brief, M2 segments of the MCA with an inner diameter (ID) ranging from 150-200 µm were meticulously dissected from brain tissues obtained from animals euthanized using 4% isoflurane. The dissected segments were then placed in ice-cold PSS_0Ca_ containing (in mM; pH 7.4): 119 NaCl, 4.7 KCl, 1.17 MgSO_4_, 18 NaHCO_3_, 5 HEPES, 1.18 NaH_2_PO_4_, 10 glucose, and 0.03 EDTA. ^6, 27, 46^

#### Assessment of Myogenic Reactivity of MCA

The impact of *Dusp5* KO on the myogenic response, myogenic tone and mechanical properties of the MCA was assessed following our published protocol. ^11, 39^ Briefly, the M2 segments of the MCA dissected and kept in ice-cold PSS_0Ca_ were mounted onto glass pipettes in a single vessel pressure myography chamber (Living System Instrumentation, Burlington, VT) that was filled with calcium-containing physiological salt solution (PSS_Ca_ = PSS_0Ca_ – EDTA + 1.6 mM CaCl_2,_ pH7.4). The chamber was connected to an IMT-2 inverted microscope (Olympus, Center Valley, PA) equipped with a digital camera (AmScope, Irvine, CA). The pH of the PSS_Ca_ was adjusted to 7.4 and maintained at 37°C using a temperature controller (Living System Instrumentation). The solution was oxygenated with a gas mixture containing 21% O_2_, 5% CO_2_, and 74% N_2_. Intraluminal pressure was regulated via a pressure servo controller (Living System Instrumentation).

The MCAs were pressurized to 40 mmHg for 30 minutes to allow the development of an equilibration phase with a spontaneous tone. The myogenic response was assessed by comparing the IDs of the vessels in response to intraluminal pressures ranging from 40 to 180 mmHg, in increments of 20 mmHg. After these experiments, the MCAs were washed with PSS_0Ca_ for six to eight times at 5 mmHg. The IDs and outer diameters were recorded.

#### Assessment of Mechanical Properties of MCA

In the formulas used in the present study, ID_0Ca_ and OD_0Ca_ represent the ID and OD measured at a specific intraluminal pressure in PSS_0Ca_, respectively; ID_Ca_ represents the ID measured at the same pressure in PSS_Ca_; and ID_0Ca5mmHg_ represents the ID measured at 5 mmHg in PSS_0Ca_. Myogenic response was evaluated by comparing inner diameters across perfusion pressures from 40 to 180 mmHg, presented as a percentage relative to the inner diameter at 40 mmHg. Myogenic tone was calculated at each transmural pressure using the previously described formula: ^40, 47^

Myogenic tone (%) = [ID_0Ca_ − ID_Ca_) / ID_0Ca_] X 100.

Wall thickness, CSA, and wall-to-lumen ratio were determined using the following formulas:

Wall thickness (µm) = (OD_0Ca_ − ID_0Ca_) / 2 CSA (µm^2^) = (Tr/4) X (OD^2^ − ID^2^)

Wall / lumen ratio = Wall thickness / ID_0Ca_

Distensibility, regarded as an essential indicator of vascular compliance, represents the ability of the artery to function as a buffer. Distensibility plays a crucial role in determining stress on the vessel wall, and a reduction in distensibility could elevate the risk of damage to the arterial wall. ^11, 39^ Incremental distensibility refers to the percentage of change in the ID_0Ca_ (ΔID_0Ca_) of the blood vessel for every 1 mmHg change in pressure (ΔP). The following equations were applied in the present study:

Distensibility (%) = (ID_0Ca_ − ID_0Ca5mmHg_) / ID_0Ca5mmHg_ X 100

Incremental distensibility (% / mmHg) = ΔID_0Ca_ / (ID_0Ca_ X ΔP) X 100

Circumferential wall strain refers to the deformation or stretching of a blood vessel, along its circumference in response to changes in pressure. Circumferential wall stress refers to the mechanical force per unit area acting on the circumferential of a blood vessel. These parameters are important in understanding the mechanical behavior of blood vessels and their ability to adapt to changes in blood flow and pressure.

Circumferential wall strain (ε) = (ID_0Ca_ – ID_0Ca 5mmHg_) / ID_0Ca5mmHg_

Circumferential wall stress (σ) = (P x ID_0Ca_) / 2WT

Elastic modulus (E = σ / ε) is an important property of evaluation of arterial stiffness. ^42^The slopes (β-values) of an exponential model with least-squares analysis of the elastic modulus (σ = σ_orig_e ^βε^; σ_orig_ is σ at 5 mmHg) were compared to determine the arterial stiffness. An increased β value (a higher value of elastic modulus) indicated a stiffer vessel.

### Statistical Analyses

All data is represented as mean values accompanied by the standard error of the mean (SEM). We employed a two-way analysis of variance (ANOVA) for repeated measures, followed by a Holm-Sidak post hoc test, to assess significant differences between *Dusp5* KO and WT rats in the pressure myograph studies. For the corresponding values of immunohistochemistry and β-values between the two groups, an unpaired t-test with Welch’s correction was utilized to determine significance. All statistical analyses were conducted using GraphPad Prism 6 software (GraphPad Software, Inc.), with statistical significance defined as *p* < 0.05.

## RESULTS

### Impacts of knockout of *Dusp5* on VSMC Content in the MCA

Immunofluorescence staining with an α-SMA antibody showed no significant difference in the number of VSMCs across the cross-section of the MCA wall between *Dusp5* KO (9,965.46 ± 465.63/mm^2^) and WT (10,884.07 ± 324.66/mm^2^) rats (Figure 1).

**Figure 1.**
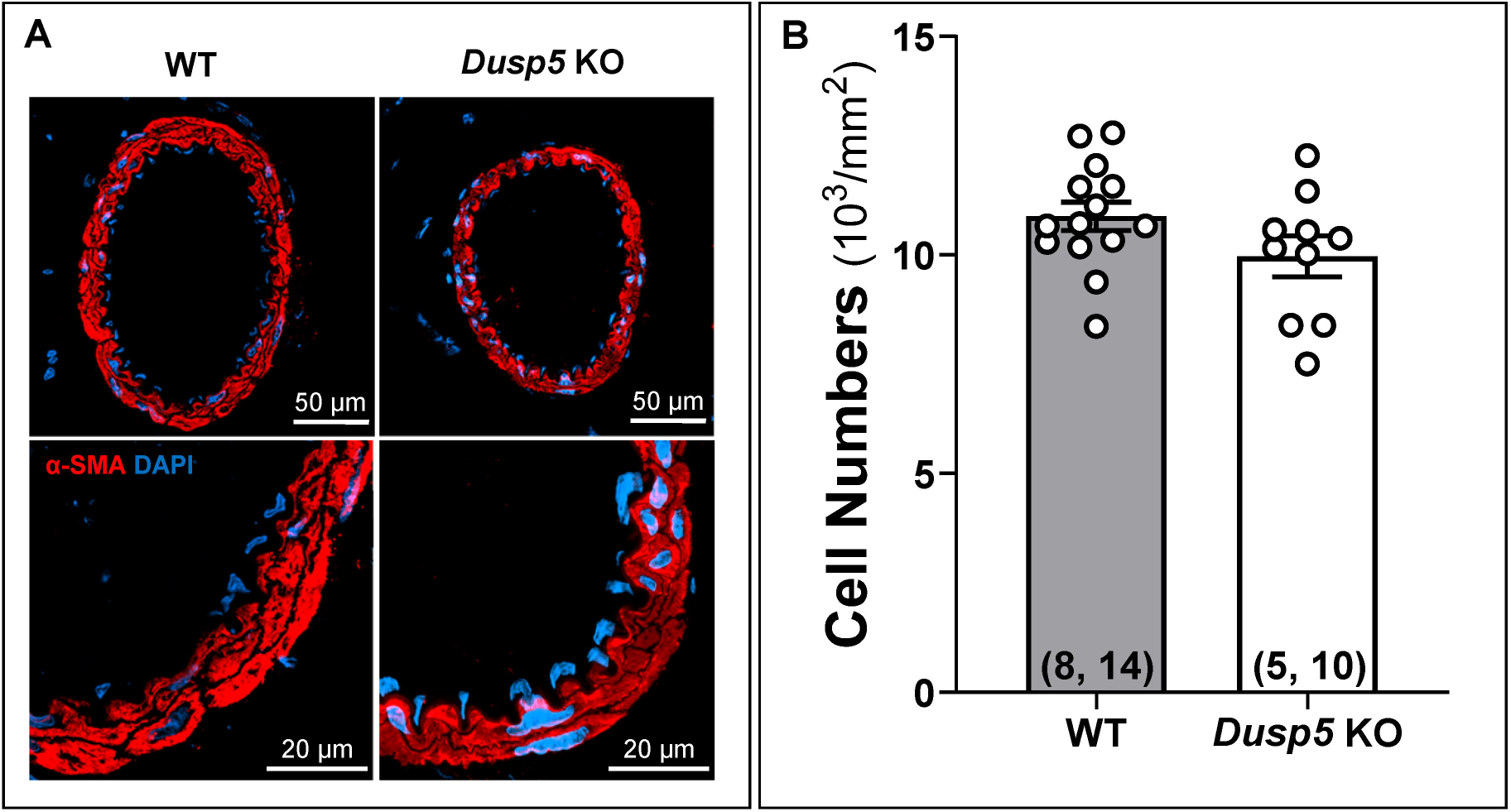
Impacts of *Dusp5* Knockout (KO) on Vascular Smooth Muscle Cell (VSMC) Content in the Middle Cerebral Artery (MCA) (A) Representative images illustrating the expression of α-SMA in the MCA wall of male *Dusp5* KO and wild-type (WT) rats. (B) Comparison of VSMC numbers across the cross-sectional area of the MCA wall in male *Dusp5* KO and WT rats. Mean values ± SEM are presented, with the numbers in parentheses indicating the number of rats and vessels studied per group (n = # rats, # vessels).

### Impacts of knockout of *Dusp5* on Collagen Content in the MCA

The differences in the collagen content between *Dusp*5 and WT rats, determined by Masson’s trichrome staining, are depicted in Figure 2. Our results indicate no significant difference in the average blue intensity, which represents the collagen-specific methyl blue, within the tunica media and tunica adventitia of the MCA between *Dusp5* KO and WT rats.

**Figure 2.**
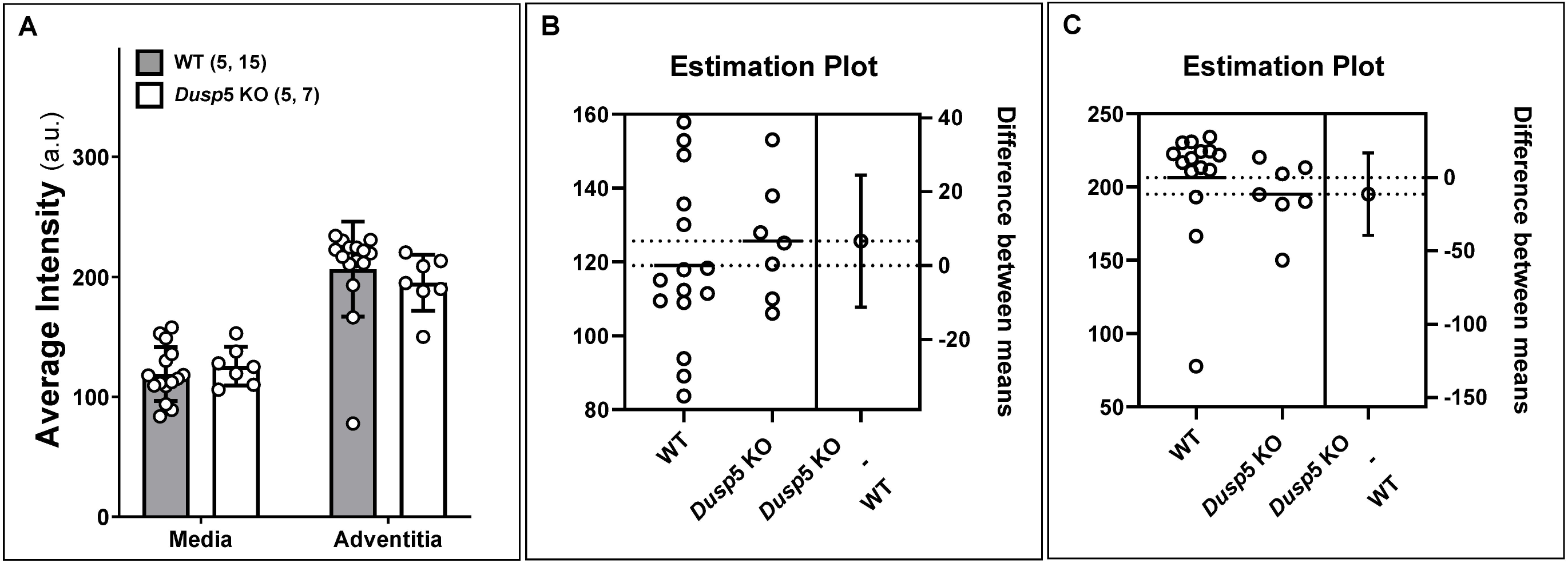
Impacts of *Dusp5* Knockout (KO) on Collagen Content in the Middle Cerebral Artery (MCA) (A) The average artificial unit (a. u.) of fluorescence intensity was compared across the cross-sectional area in the tunica media and adventitia of the MCA in male *Dusp*5 KO and wild-type (WT) rats. Estimation plots depicting fluorescence intensity in the media (B) and adventitia (C) of the MCA in *Dusp*5 KO and WT rats are presented. Mean values ± SEM are provided, with the numbers in parentheses indicating the number of rats and vessels studied per group (n = # rats, # vessels).

### Impacts of knockout of *Dusp5* on Elastin Content in the MCA

The effects of knockout of *Dusp5* on elastin content in the MCA are presented in Figure 3. We first verified the absence of the external elastic lamina in MCA in both *Dusp*5 KO and WT rats. ^39, 48^ The autofluorescence intensity was notably higher in the MCA of *Dusp*5 KO (953.76 ± 10.03 a.u.) compared to WT control rats (737.31 ± 64.88 a.u.) (Figure 3A). The thickness of the IEL was significantly increased in the MCA of *Dusp*5 KO (2.33 ± 0.02 μm) compared to WT control rats (1.85 ± 0.03 μm) (Figure 3B). Additionally, *Dusp*5 KO MCA displayed a reduced number of fenestrations and smaller fenestrae areas compared to WT controls (Figures 3C-F). These findings collectively indicate significant differences in the elastin composition and structure within the MCA wall between *Dusp*5 and WT rats.

**Figure 3.**
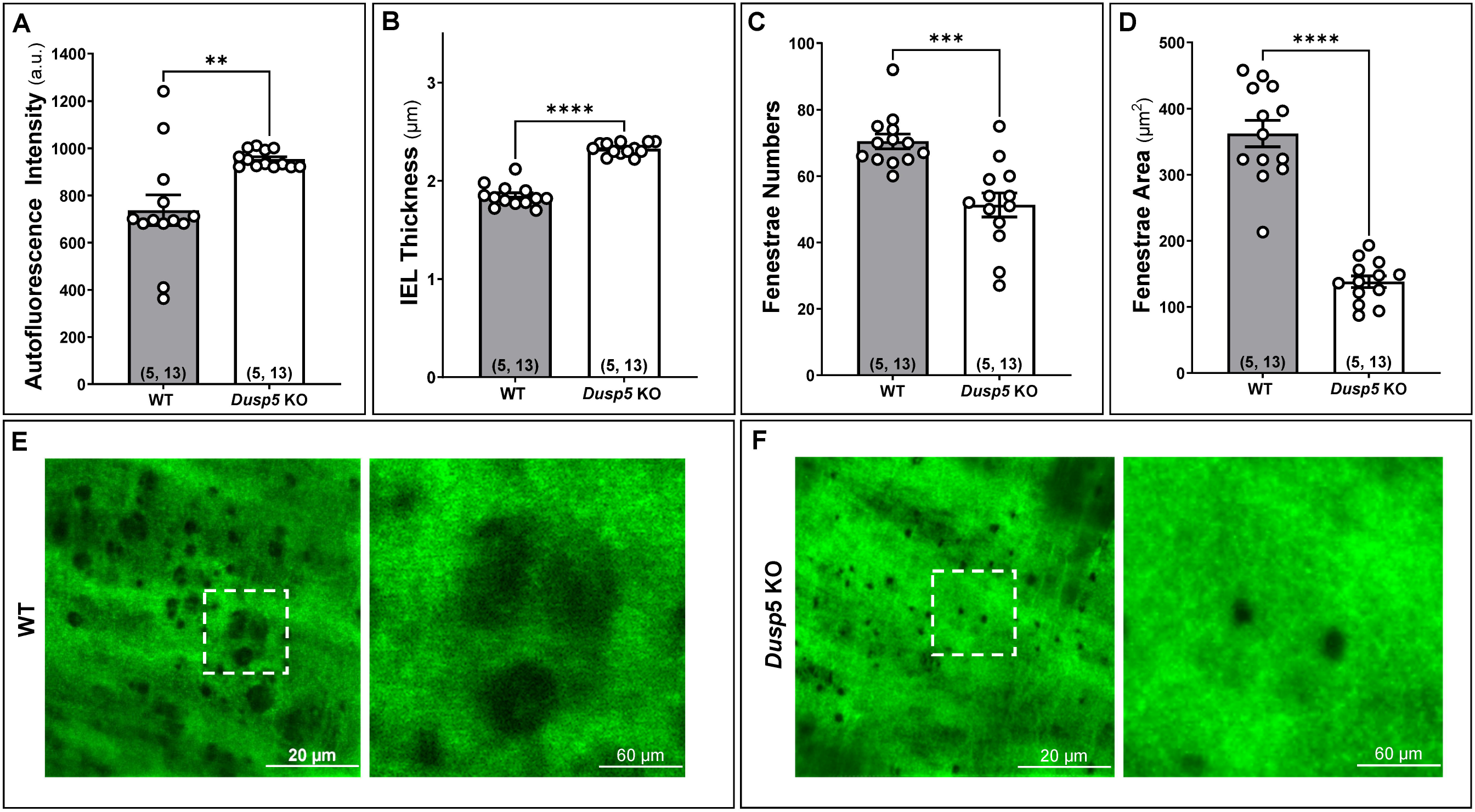
Impacts of *Dusp5* Knockout (KO) on Elastin Content in the Middle Cerebral Artery (MCA) In the MCA of male *Dusp5* KO and wild-type (WT) rats, we compared (A) the average artificial unit (a.u.) of the autofluorescence intensity in the internal elastic lamina (IEL); (B) the IEL thickness; (C) the fenestrae numbers in the IEL; (D) the fenestrae areas in the IEL. (E, F) Representative images of the elastin autofluorescence in the IEL detected by confocal microscopy. Mean values ± SEM are presented, with the numbers in parentheses indicating the number of rats and vessels studied per group (n = # rats, # vessels). One, two, three, and four asterisks (*) indicate *p* < 0.05, 0.01, 0.001, and 0.0001, respectively, from the corresponding values in *Dusp5* KO versus WT rats.

### Impacts of knockout of *Dusp5* on Myogenic Reactivity and Tone in the MCA

The baseline IDs of the MCA in *Dusp5* KO (196.79 ± 8.15 μm) and WT (205.88 ± 4.45 μm) rats were comparable. In response to increased perfusion pressure from 40 to 100, 140, and 180 mmHg, the MCA in *Dusp5* KO rats constricted to 86.89 ± 0.02%, 78.55 ± 2.02%, and 81.83 ± 2.35%, respectively (Figure 4A). These constriction responses were significantly enhanced compared to those in WT rats (91.87 ± 1.16%, 88.32 ± 2.10%, and 90.68 ± 3.10% at the same pressures), demonstrating a more dynamic and pronounced vascular response to pressure changes. On the other hand, the passive diameters (IDs_0Ca_) of the MCA were similar in *Dusp5* KO and WT rats at perfusion pressure ranging from 40 mmHg to 180 mmHg (Figure 4B). *Dusp*5 KO rats also exhibited an enhanced myogenic tone under perfusion pressures ranging from 40 to 180 mmHg (Figure 4C).

**Figure 4.**
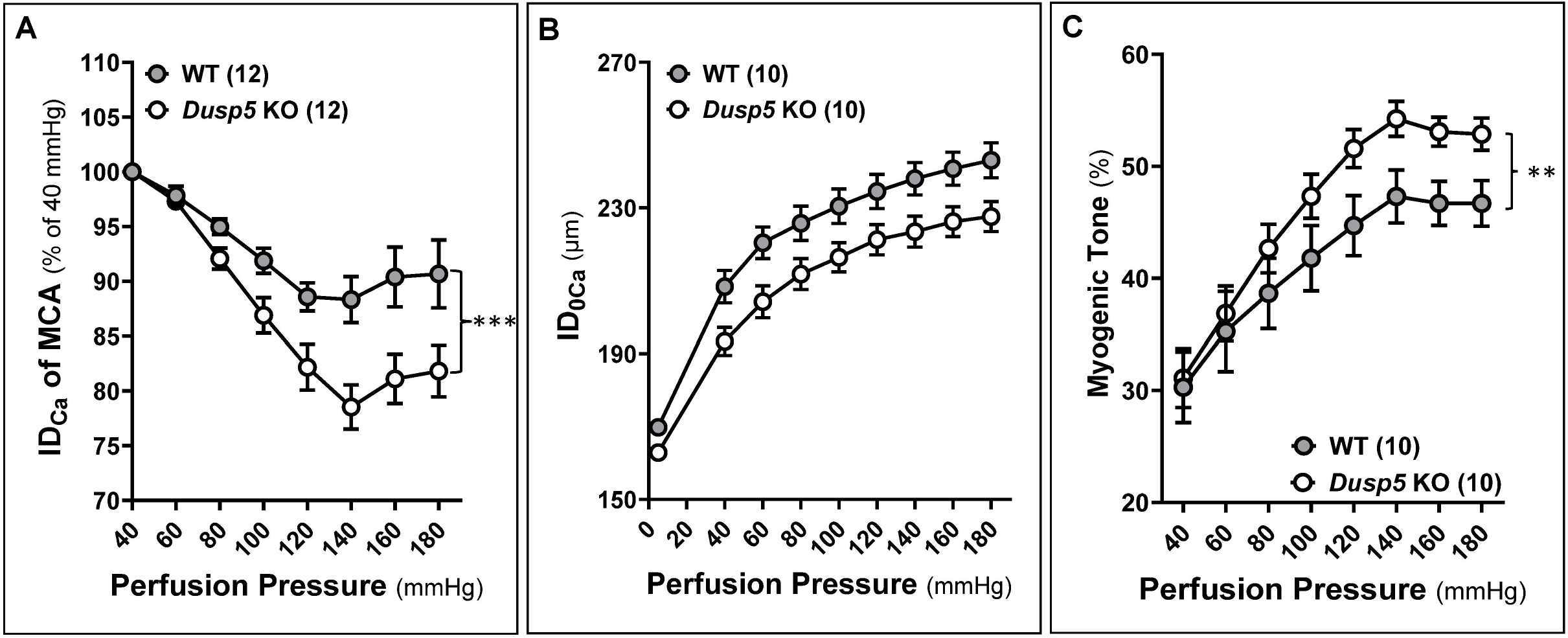
Impacts of *Dusp5* Knockout (KO) on Myogenic Reactivity and Tone in the Middle Cerebral Artery (MCA) (A) Comparison of the myogenic response of MCA of *Dusp5* KO and wild-type (WT) rats in response to an elevation in perfusion pressure ranging from 40 mmHg to 180 mmHg. (B) Comparison of the passive diameters (IDs_0Ca_) of the MCA in *Dusp5* KO and WT rats at perfusion pressure ranging from 40 mmHg to 180 mmHg. (C) Comparison of the myogenic tone of MCA of *Dusp5* KO and WT rats across perfusion pressure ranging from 40 mmHg to 180 mmHg. Mean values ± SEM are presented. Numbers in parentheses indicate the number of rats studied per group. One, two, three, and four asterisks (*) indicate adjust *p* < 0.05, 0.01, 0.001, and 0.0001, respectively, from the corresponding values in *Dusp5* KO versus WT rats after the post hoc test.

### Impacts of knockout of *Dusp5* on mechanical properties of the MCA

The wall thickness (Figure 5A), wall-to-lumen ratio (Figure 5B), and CSA (Figure 5C) of the MCA in *Dusp*5 KO rats did not show significant changes compared to wild-WT rats across perfusion pressure ranging from 40 mmHg to 180 mmHg. Moreover, the MCA in *Dusp*5 KO rats exhibits comparable distensibility (Figure 6A), incremental distensibility (Figure 6B), as well as circumferential wall stress (Figure 7A), elastic modulus (Figure 7B), and β-values (Figure 7C). These results suggest that DUSP5 is unlikely to contribute to significant changes in MCA vascular remodeling, distensibility, and stiffness under physiological conditions.

**Figure 5.**
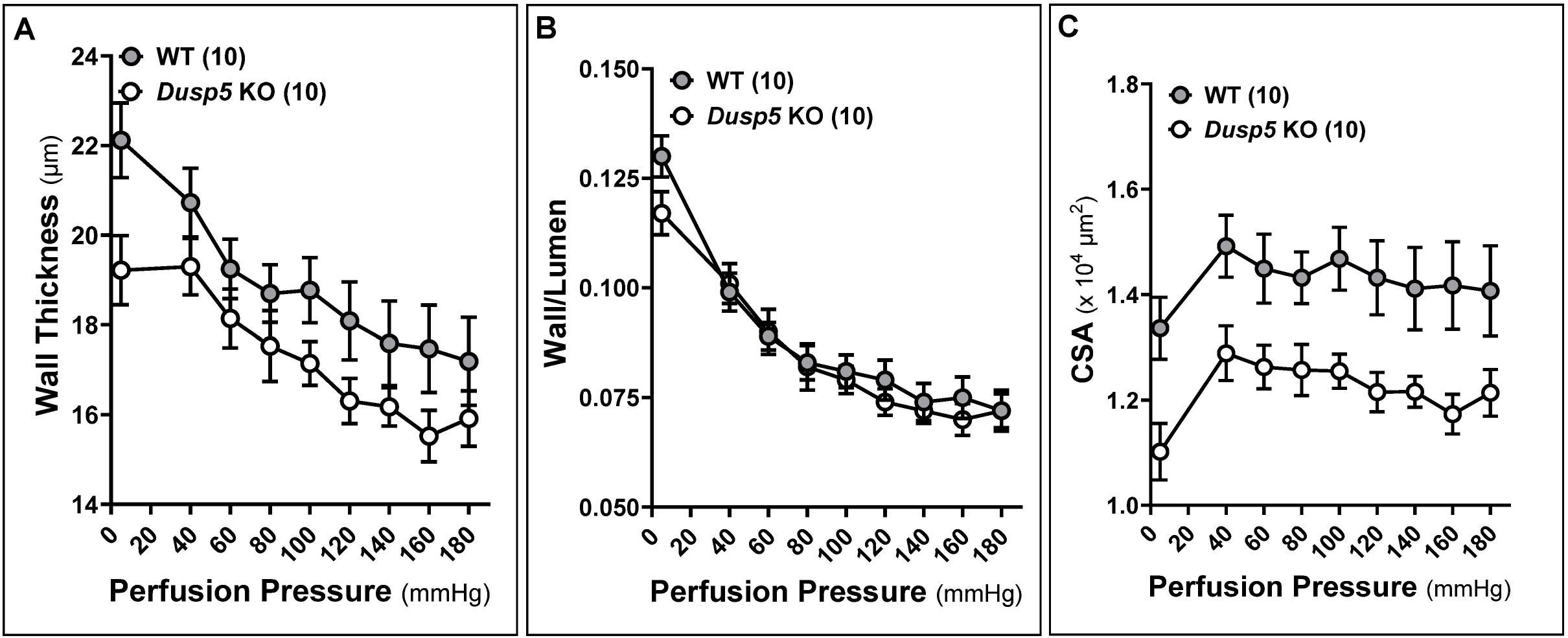
Impacts of *Dusp5* Knockout (KO) on Wall Thickness, Wall-to-Lumen Ratio, and Cross-Sectional Area (CSA) of the Middle Cerebral Artery (MCA) (A) Comparison of the wall thickness of MCA of *Dusp5* KO and wild-type (WT) rats across perfusion pressure ranging from 40 mmHg to 180 mmHg. (B) Comparison of the wall-to-lumen ratio of MCA of *Dusp5* KO and WT rats. (C) Comparison of the CSA of MCA of *Dusp5* KO and WT rats. Numbers in parentheses indicate the number of rats studied per group. One, two, three, and four asterisks (*) indicate adjust *p* < 0.05, 0.01, 0.001, and 0.0001, respectively, from the corresponding values in *Dusp5* KO versus WT rats after the post hoc test.

**Figure 6.**
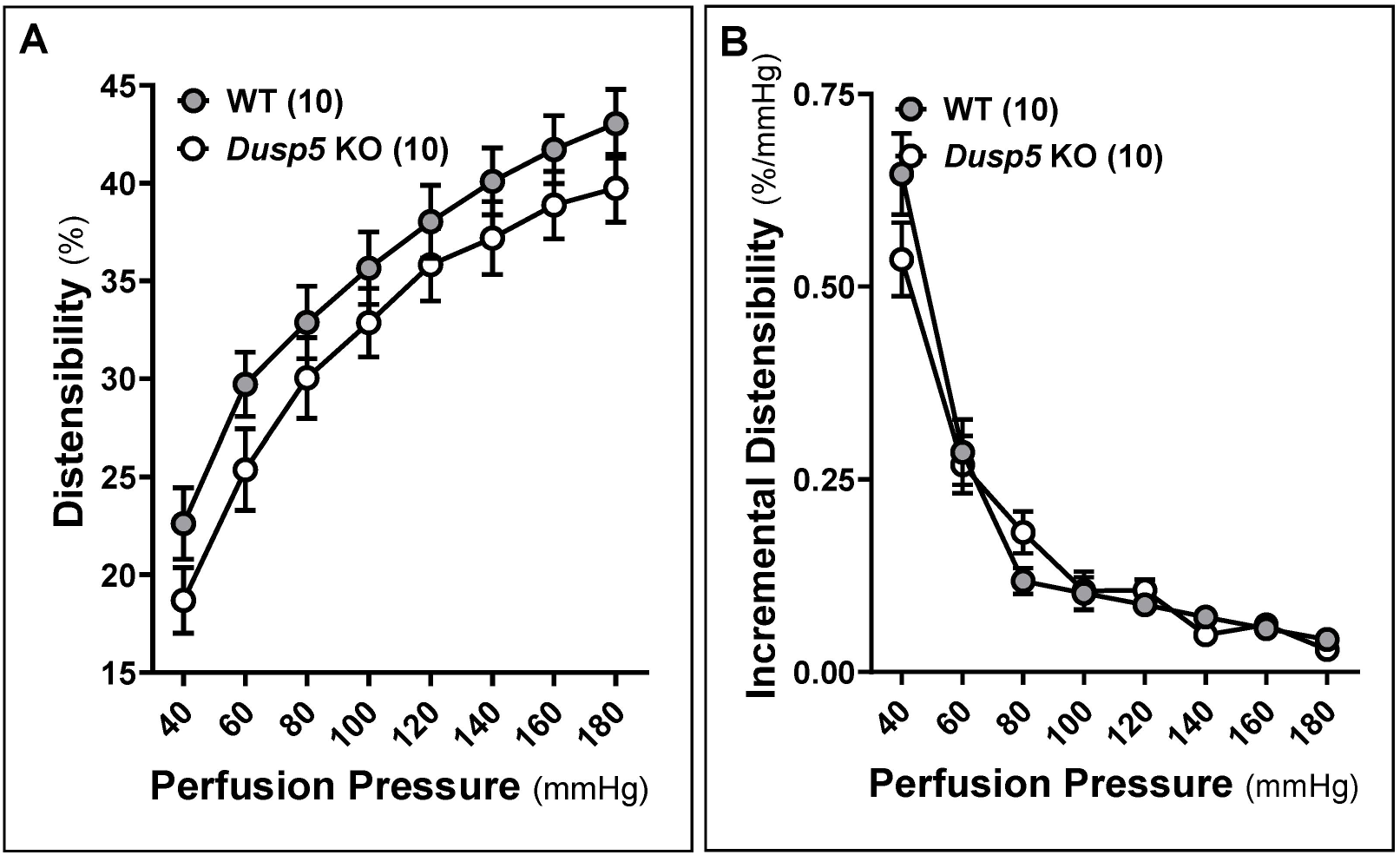
Impacts of *Dusp5* Knockout (KO) on Distensibility and Incremental Distensibility of the Middle Cerebral Artery (MCA) (A) Comparison of distensibility of MCA of *Dusp5* KO and wild-type (WT) rats across perfusion pressure ranging from 40 mmHg to 180 mmHg. (B) Comparison of incremental distensibility of MCA of *Dusp5* KO and WT rats. The numbers in parentheses indicate the number of rats studied per group. One, two, three, and four asterisks (*) indicate adjust *p* < 0.05, 0.01, 0.001, and 0.0001, respectively, from the corresponding values in *Dusp5* KO versus WT rats after the post hoc test.

**Figure 7.**
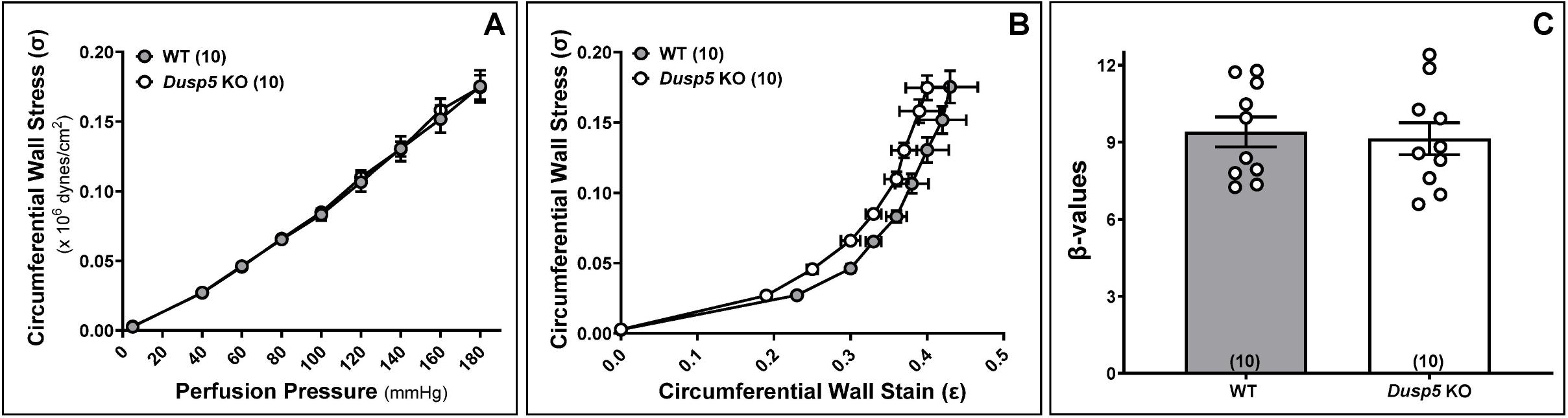
Impacts of *Dusp5* Knockout (KO) on Circumferential Wall Stress, Elastic Modulus (Stress/Strain), and β Stiffness of the Middle Cerebral Artery (MCA) (A) Comparison of circumferential wall stress of MCA of *Dusp5* KO and wild-type (WT) rats across perfusion pressure ranging from 40 mmHg to 180 mmHg. (B) Comparison of elastic Modulus (stress/strain) in MCA of *Dusp5* KO and WT rats. (C) Comparison of the slopes of the elastic modulus curves (β values) in the MCA of *Dusp5* KO and WT rats. Numbers in parentheses indicate the number of rats studied per group. 1 mmHg = 1,334 dynes/cm^2^.

## DISCUSSION

Vascular aging involves structural and functional changes in blood vessels over time, significantly impacting hemodynamics—the dynamic blood flow within the circulatory system. Aging brings about a range of vascular alterations, including changes in intrinsic vessel structure, compliance, remodeling, arterial stiffness, and endothelial function. These modifications collectively influence hemodynamic patterns, affecting the dynamic flow of blood. This, in turn, heightens the risk for various vascular diseases. ^1^ Importantly, emerging research suggests a connection between vascular aging and neurodegenerative conditions like AD/ADRD. ^49^ The compromised blood flow to the brain due to vascular aging may play a role in the development and progression of cognitive decline and neurodegeneration. ^3, 33^ Understanding the intricate relationship between vascular aging, hemodynamics, and neurological health is crucial for developing targeted interventions to mitigate the risks associated with aging and promote overall vascular and cognitive well-being.

We recently reported that KO of *Dusp*5 results in improved cerebral and renal hemodynamics and cognitive function. ^10, 11, 22, 36^ This enhancement is associated with elevated levels of pPKC and pERK1/2 in the brain and kidneys. Notably, *Dusp*5 KO also influences the passive mechanical properties of cerebral and renal arterioles, manifesting as increased myogenic tone, enhanced distensibility, greater compliance, and reduced stiffness. ^11^ These modifications in vascular properties are observed in both MCAs and PAs, coinciding with increased CBF autoregulation on the surface and in the deep cortex of the brain. While our previous studies concentrated on the PAs, the present study evaluates the impacts of *Dusp*5 KO on the structural and mechanical characteristics of the MCAs.

Intrinsic structural shifts pertain to the physical composition of vessels. In the arterial wall, the three primary layers are composed of the intima, media, and adventitia. The intima is comprised of the IEL and endothelial cells. Progressing outward, the media is characterized by elastin organized into sheets (lamellae), along with collagen, VSMCs, and the extracellular matrix. The outermost layer is formed by the adventitia, which contributes to vascular stiffness and compliance with longitudinally arranged collagen along the vessel wall. This organized structure ensures the stability, elasticity, and overall functionality of the blood vessels. ^39^ By comparing the VSMC, elastin, and collagen contents in the MCA, our study revealed no significant difference in VSMC and collagen contents between KO and WT rats. However, KO of *Dusp*5 led to notable changes in elastin content and structure in the MCA, with higher autofluorescence intensity, increased IEL thickness, and reduced elastin fenestrations compared to WT controls.

VSMCs play a pivotal role in maintaining the structural integrity of blood vessels and dynamically regulating changes in diameter in response to various stimuli. ^2, 50^ Despite the unaffected numbers of VSMCs in the context of *Dusp*5 KO, the MCA exhibits an enhanced myogenic response and increased contractility of cerebral VSMCs. ^10, 11, 36^ This suggests a potential linkage to the enzymatic function of DUSP5, implicating its role in modulating vascular dynamics and contractile properties. DUSP5 is a multifaceted regulator of the MAPK signaling pathway with broad implications in various cellular processes, vascular regulation, hypertension, renal injury, cognitive function, and cancer biology. ^12^ As an enzyme, DUSP5 exerts a negative regulatory influence on ERK1/2 in association with PKC. This interaction potentially enhances calcium influx in VSMCs and facilitates vasoconstriction in *Dusp*5 KO animals.^37^

Similar to observations in PAs, the myogenic tone in the MCA was enhanced in *Dusp*5 KO rats. Nevertheless, various passive mechanical properties of the MCA, encompassing parameters such as passive IDs, wall thickness, wall-to-lumen ratio, CSA, distensibility, incremental distensibility, circumferential wall stress, elastic modulus, and β stiffness, did not display significant alterations in the context of this study. The aforementioned parameters collectively assess vascular remodeling, compliance, distensibility, and stiffness. Our results demonstrated that KO of *Dusp*5 did not affect MCA vascular remodeling under healthy conditions. Collagen is a crucial structural component in the arterial wall, providing reinforcement and constituting the primary non-distensible element and significantly contributing to vascular stiffness and compliance. The result showed that the collagen content is comparable between *Dusp5* KO and WT rats, consistent with similar distensibility and stiffness between these two strains of animals. Elastin plays a crucial role in stabilizing the structure of arteries. When blood pressure increases, elastic fibers in the MCA wall stretch, which can trigger the myogenic response. With a thicker IEL in MCA of *Dusp*5 KO and an enhanced myogenic response, the increased tendency to constrict (in response to pressure or stretch) contributes to a higher myogenic tone. The observed differences in elastin fenestrations and IEL thickness may contribute to variations in the mechanical properties of MCA, potentially impacting its myogenic response and tone.

In conclusion, our study delves into the consequences of *Dusp*5 KO on the structural and mechanical characteristics of the MCA. Remarkably, we observe substantial increases in the thickness of the IEL and changes in elastin structure within the MCA of *Dusp*5 KO rats. Intriguingly, these alterations manifest without corresponding adjustments in VSMC numbers or collagen content in the vascular wall. The current understanding suggests that these elastic modifications, coupled with the enzymatic activity of DUSP5, likely contribute significantly to the enhanced myogenic response and tone in the MCA, as well as the reported improvements in CBF autoregulation and cognition in *Dusp*5 KO animals. Although our study exclusively focuses on young, healthy male animals, the potential impact of *Dusp*5 KO on aging vessels and vascular function-related diseases warrants further investigation.

Our study has highlighted the significant role of *Dusp*5 knockout in altering the structural and mechanical characteristics of the MCA, particularly noting the increases in the thickness of the IEL and changes in elastin structure. While these findings provide valuable insights, with limitations that are only studied in young, healthy male animals, a more detailed expanded exploration of the underlying mechanisms by which *Dusp*5 knockout influences vascular properties could be beneficial. Understanding these mechanisms in greater depth may open avenues for translating these findings into therapeutic strategies or preventive measures for vascular aging and related neurodegenerative diseases, such as AD/ADRD.

## STATEMENTS AND DECLARATIONS

### Author Contributions

HZ and FF conceived and designed research; HZ, CT, JB, YL, XF, and JRJ performed experiments; CT, HZ, JB, TJL, SB, AS, and FF analyzed data; CT, HZ, JB, RJR, and FF interpreted results; CT, HZ, JB, and FF prepared figures; CT, JB, HZ, and FF drafted the manuscript; CT, AG, CJ, TJL, SB, AS, SMS, HY, RJR, and FF edited and revised the manuscript; all authors approved the final version of the manuscript.

